# Sexual-dimorphism in human immune system aging

**DOI:** 10.1101/755702

**Authors:** Eladio J. Márquez, Cheng-han Chung, Radu Marches, Robert J. Rossi, Djamel Nehar-Belaid, Alper Eroglu, David J Mellert, George A Kuchel, Jacques Banchereau, Duygu Ucar

**Author notes:** These senior authors contributed equally to this work (George A Kuchel, Jacques Banchereau, Duygu Ucar). Sanofi US, Cambridge, MA 02139, USA.

## Abstract

Differences in immune function and responses contribute to health- and life-span disparities between sexes. However, the role of sex in immune system aging is not well understood. Here, we characterize peripheral blood mononuclear cells from 172 healthy adults 22-93 years of age using ATAC-seq, RNA-seq, and flow-cytometry. These data reveal a shared epigenomic signature of aging including declining naïve T cell and increasing monocyte/cytotoxic cell functions. These changes were greater in magnitude in men and accompanied by a male-specific genomic decline in B-cell specific loci. Age-related epigenomic changes first spike around late-thirties with similar timing and magnitude between sexes, whereas the second spike is earlier and stronger in men. Unexpectedly, genomic differences between sexes increase after age 65, with men having higher innate and pro-inflammatory activity and lower adaptive activity. Impact of age and sex on immune cell genomes can be visualized at https://immune-aging.jax.org to provide insights into future studies.

## Introduction

Human peripheral blood mononuclear cells (PBMCs) undergo both cell-intrinsic and cell-compositional changes (i.e., cell frequencies) with age, where certain immune functions are impaired and others are remodeled^1^. Analyses of human blood samples uncovered significant aging-related changes in gene expression^2^ and DNA methylation levels^3^. Recent studies revealed that chromatin accessibility of purified immune cells, especially CD8^+^ T cells, change significantly with aging, impacting the activity of important receptor molecules, signaling pathways, and transcription factors (TF)^4, 5^. Together, these changes likely contribute to aging-related immunodeficiency and ultimately to increased frequency of morbidity and mortality among older adults. However, it is unclear to what extent these aging-associated changes are shared between men and women.

Immune systems of men and women function and respond to infections and vaccination differently^6^. For example, 80% of autoimmune diseases occur in women, who typically show stronger immune responses than males^7^. Stronger responses in women produce faster pathogen clearance and better vaccine responsiveness, but also contribute to increased susceptibility to inflammatory and auto-immune diseases. Although not systematically described, these differences likely stem from differences in both cell frequencies and cell-intrinsic programs. For example, a study in young individuals showed that women have more B cells (percentage and absolute cell counts) in their blood than men^8^. Moreover, hundreds of genes <u>are</u> differentially expressed between young men and young women in sorted B cells^9^. Recently, sex-biased transcripts <u>have been</u> comprehensively described in purified immune cells (n=1,800), the majority of which <u>are</u> autosomal^10^. Furthermore, age- and sex-dependent differences in immune responses to stimuli have been observed^11, 12^. Despite the importance of sex and age in shaping immune cell functions and responses, it is not known whether men’s and women’s immune systems go through similar changes throughout their lifespan, and whether these changes occur at the same time and at the same rate.

To study this, we profiled PBMCs of healthy adults by carefully matching the ages of male and female donors. Computational pipelines for functional enrichments, trajectory and breakpoint analyses revealed immune system aging signatures including longitudinal genomic trends. These findings uncovered in which ways aging differentially affects the immune systems of men and women. These results are shared *via* a searchable R Shiny application (https://immune-aging.jax.org/).

## Results

### Profiling PBMCs of healthy adults

We recruited <u>172</u> community-dwelling healthy volunteers (<u>91 women, 81 men</u>) <u>whose ages</u> span 22-93 years old (**Figure 1a**, Supplementary **Table 1**): <u>54</u> young (ages 22-40: 23 men, <u>31</u> women), <u>59</u> middle-aged (ages 41-64: <u>31</u> men, <u>28</u> women), and <u>59</u> older subjects (65+: <u>27</u> men, <u>32</u> women). No significant differences were detected between sexes in their frailty scores or age distributions (Supplementary **Figure 1g**, Supplementary **Table 1**). PBMCs were profiled using ATAC-seq (54 men, 66 women), RNA-seq (41 men, 34 women), and flow cytometry (<u>62 men, 67 women</u>). Male and female samples for each assay were comparable in terms of frailty scores, BMI, and age except for young samples profiled with flow cytometry; young women were slightly older than men (<u>∼32.3</u> versus ∼28.35) (<u>t-test</u> p-value=<u>0.05</u>) (Supplementary **Table 1**). We took this difference into consideration while interpreting the flow cytometry data.

**Figure 1.**
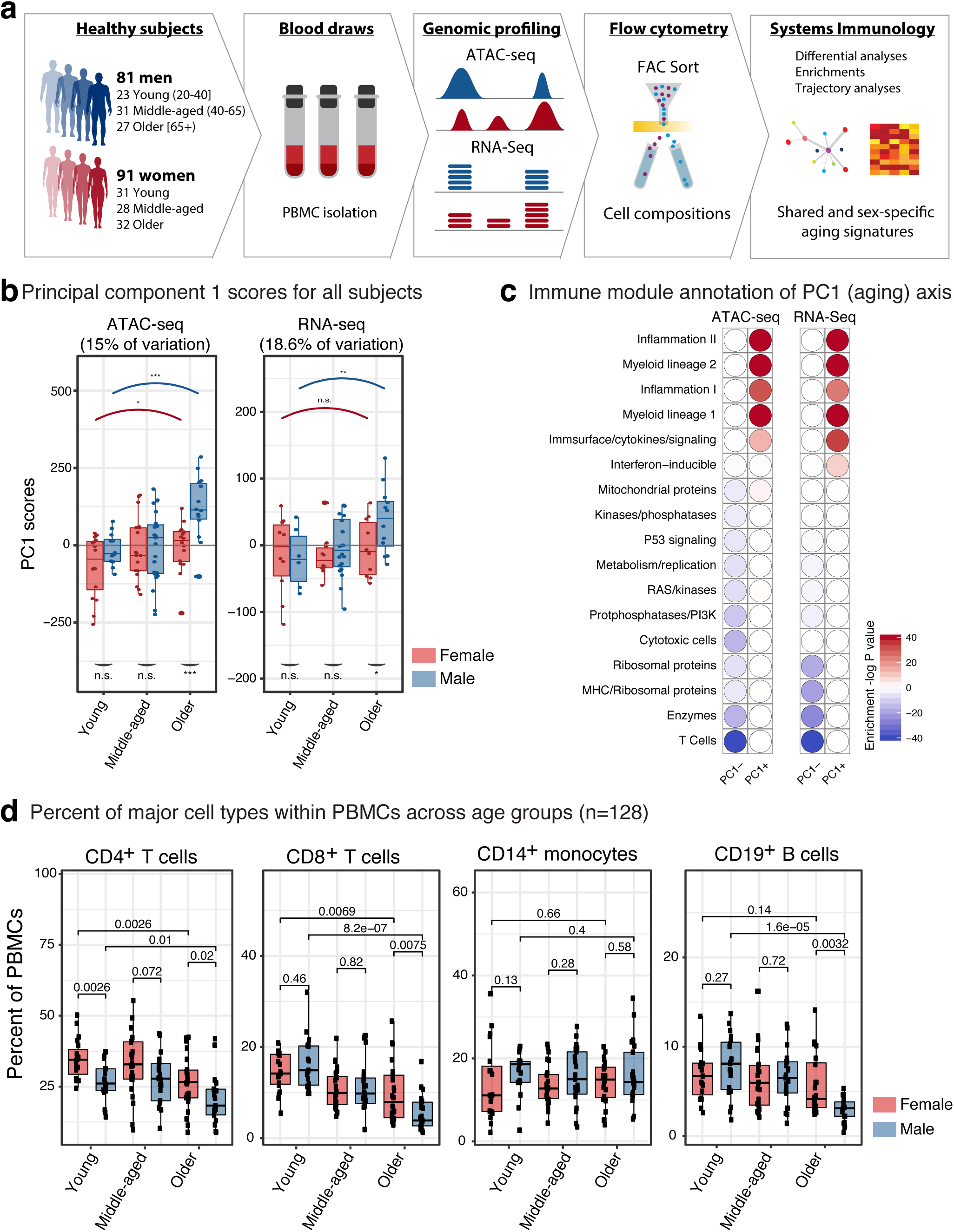
Age is the main driver of variation in PBMC <u>genomic</u> data. (**a**) Schematic summary of study design: PBMCs were isolated from blood samples of 172 healthy community-dwelling individuals (ages 22-93), from which ATAC-seq, RNA-seq and flow-cytometry data were generated. (**b**) Principal component 1 scores (PC1) were calculated for each individual from ATAC-seq (left) and RNA-seq (right) principal component analyses (PCA) results. Marker genes are selected using PC1 scores that are >= 25^th^ percentile: top positive <u>scores (ATAC-seq n=9,392 peaks, RNA-seq n=1,413 genes) and top negative scores (ATAC-seq n=10,151 peaks, RNA-seq n=1,675 genes).</u> PC1 scores from ATAC-seq and RNA-seq data increased with increasing age. Furthermore, we detected differences in PC1 scores of sexes in older subjects, where older men had higher scores than older women. (**c**) Functional enrichment of PC1-related genes based on immune modules^22^. Note that myeloid/inflammation related genes were associated with high and positive scores, whereas adaptive immunity/lymphoid related genes were associated with high and negative PC1 scores. These enrichments align well with an age-related increase in the myeloid lineage and inflammation, and an age-related decline in naïve T cell activity. Hypergeometric test was used to calculate enrichment p-values. (**d**) Flow cytometry data from <u>128</u> individuals that reflect proportions of major cell types within PBMCs. Subjects were stratified based on age group and sex: Young: 22-40, Middle-aged: 41-64, Older: 65+ years old. <u>Wilcoxon rank-sum test was used to compare data from female (n=67) and male (n=61) subjects.</u> Note that T cell proportions decline with age in both sexes, whereas the decline in B cell proportions is specific to older men. <u>Box plots represent median and IQR values, whiskers extend to 1.5 times the IQR; ***p<0.001, **p<0.01, *p<0.05, n.s.: non-significant.</u> Source data are provided as a Source Data file for panel c.

### Aging is the main driver of variation in PBMC ATAC-seq and RNA-seq data

To identify major sources of variation in genomics data, we conducted principal component analyses (PCA) using open chromatin regions (OCRs; n=78,167) and expressed genes (n=12,350) from high quality samples (100 ATAC-seq, 74 RNA-seq) (Supplementary **Table 2**). The first principal component (PC1) captured 15% of the variation in ATAC-seq and 18.6% in RNA-seq data and associated to age groups (**Figure 1b**, Supplementary Figures **1a, 1b**, Supplementary **Table 3**). PC1 differences between young and old samples were more significant in men (ATAC-seq: p<0.05 versus p<0.001, RNA-seq: p=N.S. versus p=0.03, <u>Wilcoxon rank-sum test</u>) (**Figure 1b**). Annotation of PC1-related genes using immune modules^13^ revealed genomic declines associated with the ‘T cell’ module and gains associated with ‘myeloid lineage’ and ‘inflammation’ (**Figure 1c**), confirmed by enrichment analyses based on single-cell RNA-seq data (Supplementary **Figure 1c**). Together, these results suggest that aging is the main driver of variation in PBMC epigenomes/transcriptomes, where age-related variation is negatively associated with naïve T cells and positively associated with myeloid lineage and inflammation, in agreement with the signatures of immunosenescence including impaired responsiveness of adaptive immunity and increases in low-grade and systemic inflammation (i.e., inflamm-aging) with age, which together contribute to aging diseases^14^.

### Aging-related changes in PBMC cell compositions

We quantified the frequencies of CD4^+^ and CD8^+^ T cells, CD19^+^ B cells, and CD14^+^ monocytes in PBMCs (67 women, 62 men) (Supplementary **Table 4**, <u>Supplementary Figure 2a</u>). Among these, the proportions of CD8^+^ T cells were the most significantly affected with age (p=8.2e^-07^ for men, p=0.0069 for women, <u>Wilcoxon</u>) (**Figure 1d**). Similarly, CD4^+^ T cell frequencies declined both in men (<u>Wilcoxon</u> p=0.01) and women (<u>Wilcoxon</u> p=<u>0.0026</u>). However, we detected a male-specific decline in B cell proportions after age 65 (<u>Wilcoxon</u> p=1.6e^-05^), resulting in a significant difference in B cell proportions between older men and older women (<u>Wilcoxon</u> p=0.0032) (**Figure 1d**, Supplementary **Table 4**). We did not detect significant aging- or sex-related changes in CD14^+^ monocytes (Kruskal-Wallis p-value=<u>0.32</u>) (**Figure 1d**). These four cell types constitute the majority of PBMCs. However, PBMCs also include NK cells (5%), DCs (1-2%), CD16^+^ monocytes (1%), ILCs (<1%)^15^. Among these, the abundance of CD16^+^ monocytes reported to increase with age, whereas ILC and DC subsets and CD56^+^ NK cells decrease with age^16^. Furthermore, CD69^+^ CD16^+^ NK cells were more abundant in men, whereas subsets of T cells and ILCs showed more abundance in women.

**Figure 2.**
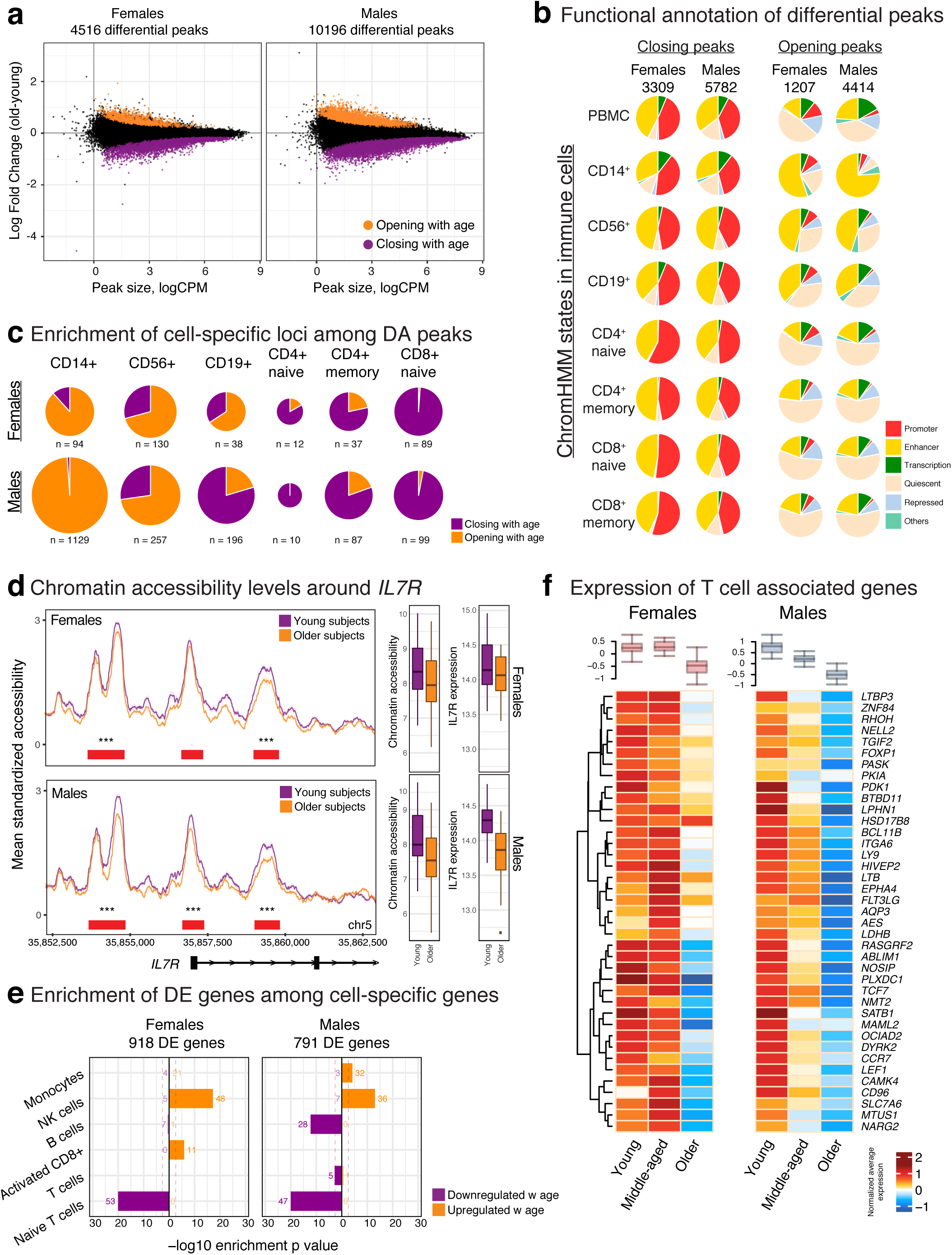
Shared and sex-specific <u>epigenomic</u> signatures of aging. (**a**) <u>Fold change (older-young) versus read counts per million (logCPM) from ATAC-seq differential analyses in women (left) and men (right) based on 78,167 ATAC-seq peaks (5% FDR after Benjamini-Hochberg P-value adjustment, 58 subjects).</u> Differentially opening/closing peaks are represented in orange/purple. (**b**) Functional annotations of differentially accessible (DA) peaks in men and women using chromHMM states in PBMCs and cell subsets. Closing peaks tend to be enhancers/promoters across all cell types, whereas opening peaks tend to be enhancers in monocytes and NK cells. (**c**) Distribution of DA peaks among cell-specific loci from chromHMM states. Innate cell-specific loci (monocytes, NK cells) are enriched in opening peaks, T cell-specific loci are enriched in closing peaks. B cell-specific loci are more likely to be an opening peak in women and closing peak in men. (**d**) Left: Average chromatin accessibility profiles (grouped by age and sex) around the *IL7R* locus - is associated with chromatin closing with age- in women (top<u>, n=32</u>) and men (bottom<u>, n=26</u>). <u>Red bars indicate peaks and their significance value from differential analyses (*p<0.05, **p<0.01, ***p<0.001, no symbols: non-significant).</u> Right: Normalized chromatin accessibility (<u>combined from</u> three peaks shown in the figure) and gene expression levels for *IL7R* in young <u>(n=16)</u> and older <u>(n=16)</u> women (top) and <u>young (n=12) and older (n=14)</u> men (bottom). In both sexes, chromatin accessibility and gene expression levels decline with age albeit more significantly in men. (**e**) Enrichment of differentially expressed (DE) genes using cell-specific gene sets from scRNA-seq data. Enrichment p-values are calculated using hypergeometric test. Gene expression programs for innate cells (NK, monocytes) are activated, whereas programs for T cells are inactivated in both sexes. B-cell specific genes were downregulated specifically in men. (**f**) Average expression levels of T cell-specific genes grouped by age group and sex. Note the decline in both sexes. Box plots on the top summarize the data from all genes represented in the heatmap. <u>All box plots represent median and IQR values, with whiskers extending to 1.5 times the IQR.</u> Source data are provided for panels a,b,c,e.

We studied subsets of CD4^+^ and CD8^+^ T cells in a subgroup of young and older individuals (n=<u>80</u>): naïve (CD45RA^+^, CCR7^+^), central memory (CD45RA^-^, CCR7^+^), effector memory (CD45RA^-^, CCR7^-^), and effector memory RA (CD45RA^+^, CCR7^-^). Expectedly, naïve T cell frequencies decreased with age, particularly in CD8^+^ T cells (Supplementary **Figures 1d, 1e**). As previously reported^17^, women had more naïve T cells compared to men. For example, <u>∼13.2%</u> of PBMCs were naïve CD4^+^ T cells in young women, compared to ∼7.8% in young men (<u>Wilcoxon</u> p=<u>0.00057</u>) despite the cohort of young women being slightly older (Supplementary **Tables 1, 4**). Women have elevated thymic function compared to men at all ages^18^, potentially explaining observed sex-differences in naïve T cells. Sex- and age-related changes in other T cell subsets were not as significant and as consistent as the changes in naïve T cells (Supplementary **Figures 1d, 1e**).

Using generalized linear models (GLM), we studied the relationship between age and sex of individuals and the variation in their PBMC compositions, which confirmed that age is most strongly associated with CD8^+^ T cell proportions, especially for naïve cells (Supplementary **Figure 1f**). Whereas sex is most strongly associated with naïve CD4^+^ T cells; women have higher proportions compared to men (Supplementary **Figures 1d, f**). Finally, we observed that B cell proportions are associated with joint effects of sex and age (Supplementary **Figure 1f**), due to the male-specific decline in B cell proportions after age 65 (**Figure 1d**). Together, these data uncovered shared and sex-specific changes in PBMC cell compositions with age, some of which underlie genomic changes.

### Shared and sex-specific chromatin accessibility signatures of aging

We detected ATAC-seq peaks that are differentially accessible (DA) between young and older individuals^19^ (FDR 5%). In men, 10,196 peaks were DA with age (13% of tested: 5,782 closing, 4,414 opening), compared to 4,516 DA peaks in women (5.8% of tested: 3,309 closing, 1,207 opening) (**Figure 2a**, Supplementary **Table 5**). Male and female DA peaks significantly overlapped, despite thousands of sex-specific loci associated with aging. Furthermore, a significant correlation was detected between sexes in terms of their epigenomic aging signatures (<u>Pearson</u> *r*=0.445, p<0.001). DA peaks corresponded to 6,612 and 3,634 genes in men and women, respectively (Supplementary **Figure** 2b). Functional annotation of these peaks using Roadmap chromatin states in immune cells^20^ revealed that closing peaks were mostly regulatory elements (promoters, enhancers) shared across cells, especially T cell subsets (**Figure 2b**, Supplementary **Figure** 2c). In contrast, opening peaks were mostly enhancers of CD14^+^ monocytes and CD56^+^ NK cells in both sexes (**Figure 2b**, Supplementary **Figure** 2d). As previously reported^5^, there were more ‘quiescent’ loci (i.e., loci without functional annotation) among opening peaks, which may be due to global chromatin deregulation with age^21^.

To identify immune cells/functions affected by aging, we studied changes at cell-specific loci (i.e., loci active in one cell type) (Supplementary **Figure** 2c). In both sexes, opening peaks were associated with monocyte-, NK, and memory CD8^+^ cell-specific loci, whereas closing peaks were associated with naïve T cell-specific loci (**Figure 2c**, Supplementary **Figures** 2e, 2f). Despite these similarities, a greater number of cell-specific loci were affected with aging in men. Notably, 960 monocyte-specific peaks significantly gained accessibility with aging in men, whereas only 64 such loci were detected in women. Only B cells showed the opposite changes between sexes, where B cell-specific loci were more likely to be among opening peaks in women and among closing peaks in men (**Figure 2c**). Fold change distributions and enrichment analyses confirmed these results. Interestingly, closing peaks were enriched in B cell-specific loci only in men (Supplementary **Figures** 2e, 2f). Immune module^13, 22^ and WikiPathways enrichment analyses uncovered that inflammation related modules/pathways were associated to chromatin opening in both sexes (Supplementary **Table 6**) (e.g., MyD88 cascade (WP2801) pathway). In contrast, closing peaks were significantly associated to modules/pathways related to T cells, notably the IL-7 Signaling pathway as we previously observed^5^. Several molecules in this pathway were associated to chromatin closing in both sexes including *IL7R, JAK1, STAT5B* and *PTK2B* (Supplementary **Table 6**, Supplementary **Figure 3** for more examples). Together, these data uncovered an epigenomic signature of aging shared between sexes, which include gains in chromatin accessibility for pro-inflammatory processes, monocytes and cytotoxic cells (NK, CD8^+^ memory) and losses in accessibility for naïve T cells. Interestingly, these changes were more pronounced in men, despite cohorts being comparable for age, frailty, and BMI (Supplementary **Figure 1g**, Supplementary **Table 1**). Furthermore, we discovered that B cells age differently between sexes, where a significant loss in chromatin accessibility was detected only in men.

**Figure 3.**
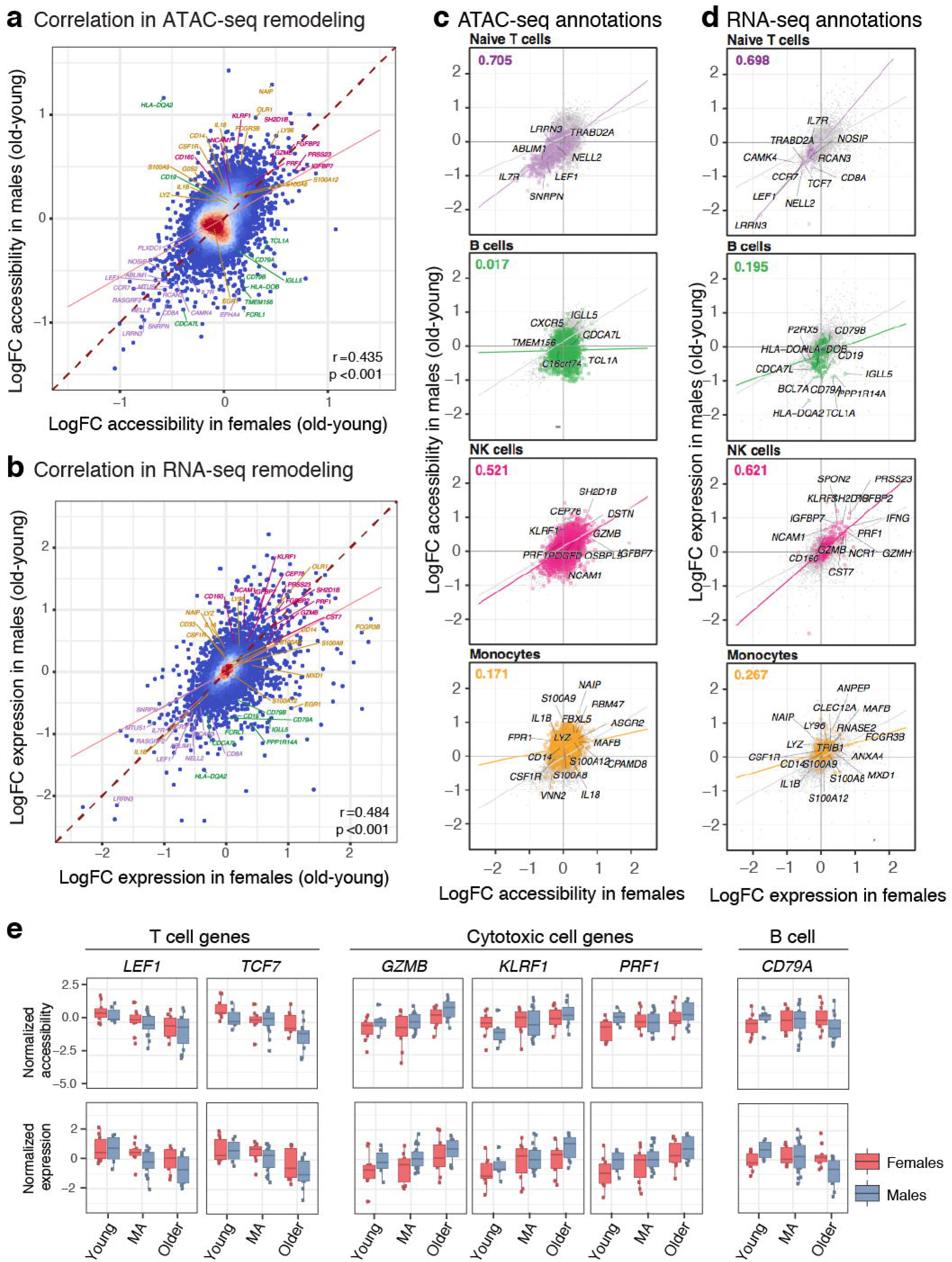
<u>Sex-dimorphic changes in monocyte- and B cell-associated loci</u>. Correlation of age-related ATAC-seq (**a**) and RNA-seq (**b**) remodeling between women and men PBMCs. Note the overall <u>large and positive Pearson correlation coefficients</u>. <u>Genes are associated to ATAC-seq peaks based on nearest TSS, and are color coded in both plots according to their association to immune modules</u> (purple: T cells, green: B cells, pink: NK cells, yellow: monocytes). <u>Only regulatory (TSS/enhancer) peaks are included and peaks-genes are matched between both plots (n=10,707 loci). Blue-red gradient</u> on data points represents their <u>relative local</u> density. (**c**) Correlation between sexes for age-related ATAC-seq remodeling stratified by cell-specific loci from chromHMM annotations. Note that the highest correlation is observed in naïve T cells <u>(n=833 peaks)</u>, which is associated with negative fold changes (i.e., loss in chromatin accessibility with age) in both sexes. Chromatin remodeling correlates the least between sexes for B <u>cell- (n=1,645 peaks)</u> and monocyte-specific loci <u>(n=6861 peaks),</u> with NK cells (n=3,008 peaks) showing a positive trend in both sexes. (**d**) Correlation between sexes for age-related RNA-seq remodeling stratified by cell-specific genes from single-cell RNA-seq data. Note that the highest correlation is observed in naïve T cells <u>(n=70 genes)</u>, which is associated with negative fold changes (i.e., downregulation with age) in both sexes. On the other hand, NK cells <u>(n=403 genes)</u> are highly correlated between sexes and associated with increased expression with age in both sexes. In agreement with ATAC-seq data, gene expression remodeling correlates the least between sexes for B <u>cell- (n=144)</u> and monocyte-specific genes <u>(n=748)</u>. Values at the top left corner represent the <u>Pearson</u> correlation coefficient between sexes only for the genes/loci associated to that cell type. (**e**) Normalized expression and accessibility levels for important molecules associated to T, cytotoxic, and B cells. <u>LogFC = mean log2 of fold changes between old and young samples;</u> MA = middle-aged. Box plots represent median <u>and IQR values, with whiskers extending to 1.5 times the IQR.</u> Source data are provided as a Source Data file for panels a,b,c,d.

### Correlated aging-related changes in transcriptomes and epigenomes

From PBMC RNA-seq data, we identified 918 differentially expressed (DE) genes in women (539 up, 379 down) and 791 genes in men (510 up, 281 down) (FDR 10%)^19^ (Supplementary **Figure 4a**, Supplementary **Table 7**). DE genes overlapped significantly between sexes. For example, 201 downregulated genes were shared (<u>Chi-square</u> p-value=8.12e-149) and included important T cell molecules (e.g., *TCF7, LEF1, CCR7*) (Supplementary **Figure 4b**). Annotations using cell-specific marker genes from PBMC scRNA-seq data and sorted cell transcriptomes^10^, revealed that upregulated genes were most significantly enriched in marker genes for NK (**Figure 2e**), monocytes and activated CD8^+^ T cells (Supplementary **Figures 4c, 4d**, Supplementary **Table 8**). In contrast, downregulated genes were most significantly enriched in naïve T cell marker genes (e.g., *CD27, CCR7, LEF1*) in both sexes (**Figures 2e, 2f**, Supplementary **Table 8**). Similar to the ATAC-seq data (**Figure 2c**), aging-related gene expression changes in the B cell compartment were sex dimorphic. B cell-specific genes, especially naïve B cells (e.g., *BCL7A, PAX5, CD79A*) were downregulated with age only in men (**Figure 2e**, Supplementary **Figure 4d, 4e**), potentially due to changes in cell composition (**Figure 1d**). Notably, aging-related transcriptomic and epigenomic changes correlated significantly in men (<u>Pearson</u> *r*=0.31, p<10e^-4^) and in women (<u>Pearson</u> *r*=0.21, p<10e^-4^). Enrichment analyses confirmed the upregulation of genes associated to cytotoxic cells and inflammation in both sexes (e.g., *GZMB, PRF1, NKG7* in women) (Supplementary **Figure 4f**) and downregulation of T cell genes (e.g., *LEF1, TCF7, CCR7* in both sexes) (Supplementary **Table 8**). These results demonstrate that age-related changes in epigenomes and transcriptomes correlated significantly and uncovered an age-related shift in PBMCs from adaptive to innate immunity in both sexes, albeit more pronounced in men.

**Figure 4.**
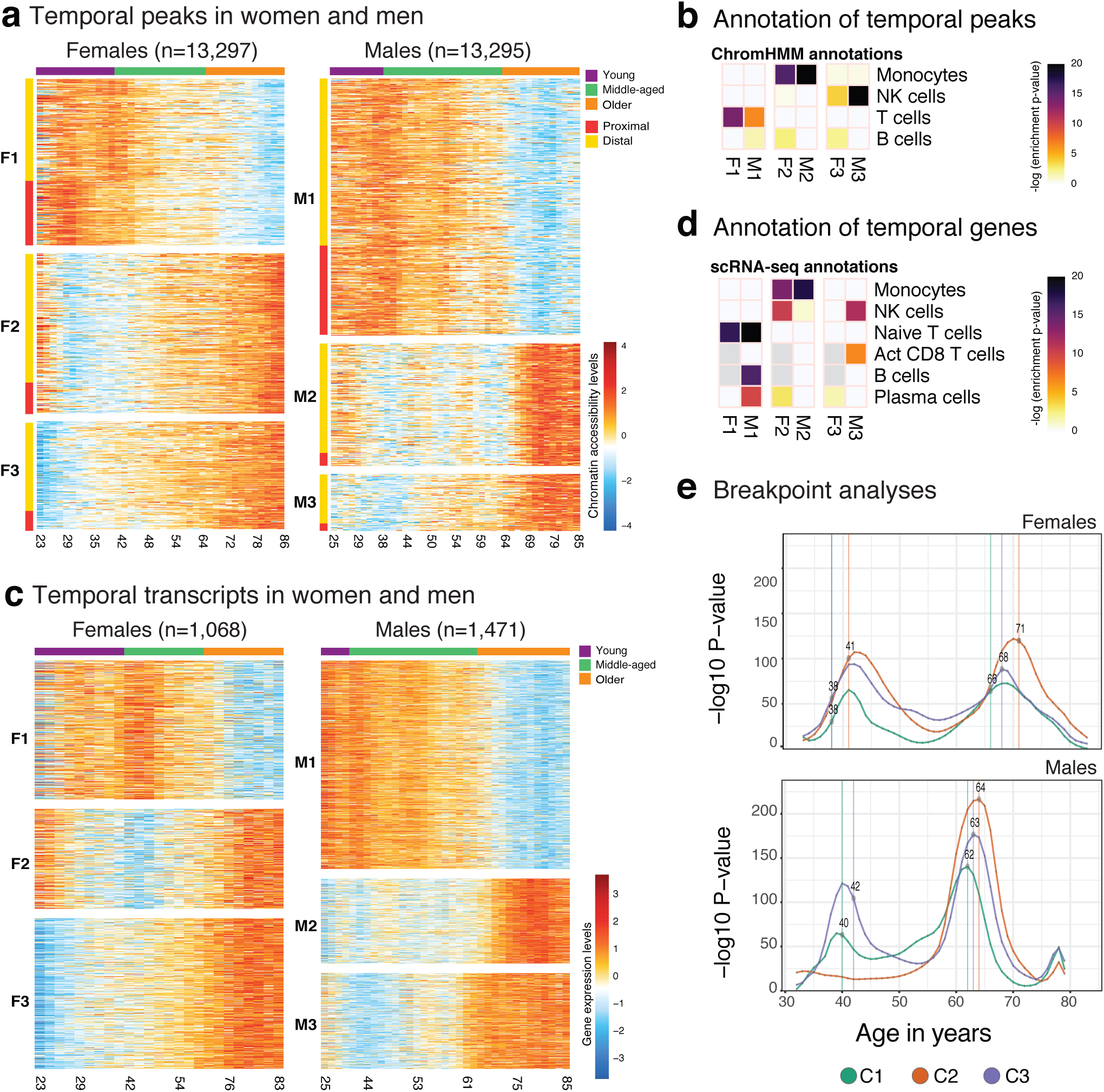
<u>Epigenome and transcriptome</u> changes over human adult lifespan. (**a**) Heatmap of ATAC-seq data (fitted values from ARIMA models) with a significant chronological trend in women (left, n=13,297) and men (right, n=13,295), <u>as a function of age in years</u>. Values represent z-score normalized accessibility values relative to the row <u>(i.e., peak)</u> mean. K-means clustering was used to group these peaks into three clusters in men and women (M1/F1, M2/F2, M3/F3). Color bar on the top represents discrete age groupings as defined in this study (young, middle-aged, older). Rows are annotated according to their position relative to the nearest TSS: proximal if <1kbp distance, distal <u>otherwise</u>. (**b**) ChromHMM state annotations of temporal peak clusters in women (F1-F3) and men (M1-M3). Colors represent hypergeometric enrichment test p-values; <u>light grey cells indicate insufficient number of genes to run an enrichment test.</u> (**c**) Heatmap of ARIMA-fitted expression values for genes with a significant chronological trend in women (left, n=1,068) and men (right, n=1,471), <u>as a function of age in years</u>. Three clusters per sex are identified using k-means clustering: F1-3 and M1-3. Values represent z-score normalized expression values relative to the row <u>(i.e., gene)</u> mean. (**d**) Annotation of temporal gene clusters using cell-specific gene sets derived from single-cell RNA-seq data. <u>Colors represent hypergeometric enrichment test p-values; light grey cells in plot indicate that there were insufficient genes in the gene sets to run an enrichment test.</u> (**e**) <u>Inverse log p-</u>value distributions from breakpoint analysis for each cluster for women (top) and men (bottom), where curve height indicates magnitude of differences between preceding and succeeding age windows. Note that there are two age brackets where epigenomic changes take place abruptly both in men and women. Points and vertical lines mark median age estimates for a breakpoint integrated over multiple scales, <u>and C1-C3 correspond to temporal clusters M1-M3 or F1-F3 from each sex</u> (see Methods and **Figure S5c** for details). Source data are provided as a Source Data file for panels a,b,c,d,e.

### Age-related changes in monocyte- and B cell-associated loci differ between sexes

Age-related changes in ATAC-seq (**Figure 3a**, <u>Pearson</u> *r*=0.435, *p*<0.001) and RNA-seq data (**Figure 3b**, <u>Pearson</u> *r*=0.484, *p*<0.001) positively correlated between sexes. However, enrichment analyses using Roadmap^20^ and scRNA-seq data revealed that similarity between sexes depends on the cell type (**Figures 3c, 3d**). For example, decreases associated with naïve T cells were highly correlated between sexes both in transcriptomic (<u>Pearson</u> *r*=0.698, **Figure 3d**) and epigenomic data (<u>Pearson</u> *r*=0.705, **Figure 3c**, Supplementary **Figure 5a**), likely due to the age-related loss of naïve T cells (Supplementary **Figures 1d, 1e**). These included shared changes observed in *IL7R* – a receptor associated to aging (**Figure 2d**)^5^ - chemokine receptor *CCR7*, co-receptor *CD8A*, and TFs *TCF7* and *LEF1* (Supplementary **Table 6**). Gene expression levels of these molecules also decreased with age in both sexes (**Figures 2d, 3e**). Similarly, changes in cytotoxic cells were highly correlated between sexes (<u>Pearson coefficient</u> NK cells: RNA-seq *r*=0.621, ATAC-seq *r*=0.521; cytotoxicity module: RNA-seq *r*=0.772, ATAC-seq *r*=0.675) (**Figures 3c, 3d**, Supplementary **Figures 5b, 5c**) and their activity increased with age in both sexes. These included granzyme B (*GZMB*) and perforin (*PRF1*), which have complementary cytotoxic functions, and *KLRF1*, which is selectively expressed in NK cells^23^ relative to cytotoxic T cells (CTLs) (**Figure 3e**). Combined with our previous study^5^, these results suggest that both CTLs and NK cells contribute to the increased cytotoxicity with age in both sexes. In contrast, age-related changes associated to monocytes and B cells showed the lowest correlation between sexes (**Figures 3c, 3d**, Supplementary **Figures 5b, 5c**). In monocytes, a majority of peaks/genes were activated with age in both sexes; however, the magnitude of activation was more pronounced in men (**Figure 2c**), which reduced the correlations (<u>Pearson coefficient</u> RNA-seq *r*=0.267, ATAC-seq *r*=0.171) (**Figures 3c, 3d**). These included pro-inflammatory cytokines *IL18, IL1B* and *S100A8/S100A9* genes that modulate inflammatory responses and serve as potential biomarkers of inflammation-related diseases^24^. In B cells, the distinction between sexes stemmed from the male-specific downregulation/chromatin closing of B-cell specific loci/genes (<u>Pearson coefficient</u> RNA-seq *r*=0.195, ATAC-seq *r*=0.017) (**Figures 3c, 3d**), including the chemokine receptor *CXCR4* and B cell receptor *CD79A* (**Figure 3e**, Supplementary **Table 6**).

**Figure 5.**
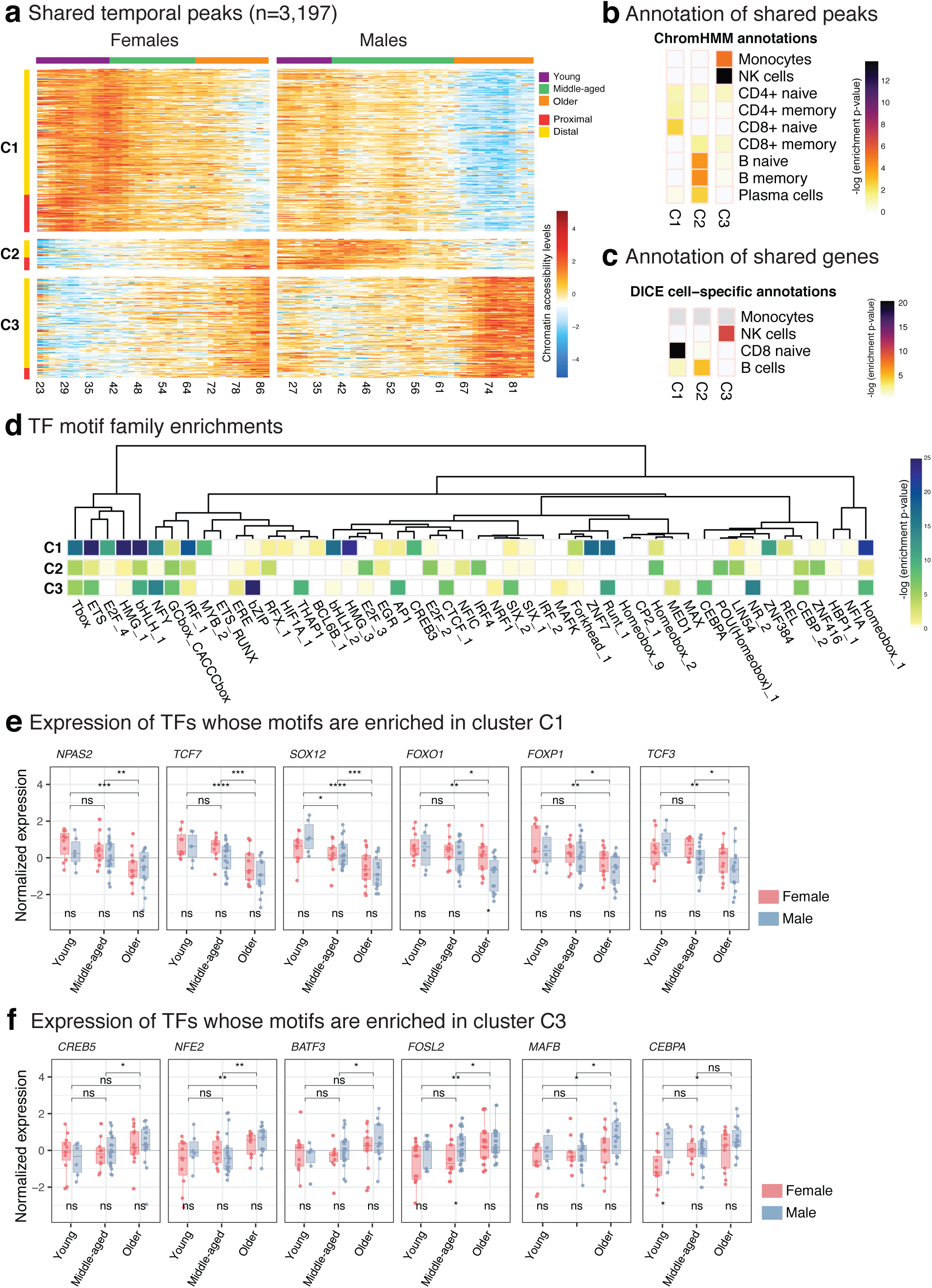
<u>Sex-specific patterns in temporal peaks/genes</u>. **(a)** Heatmap of ATAC-seq peaks (fitted values from ARIMA models) with a chronological trend in both women and men (n=3,197), <u>as a function of age in years</u>. Values represent z-score normalized accessibility values relative to the row <u>(i.e., peak)</u> mean. K-means clustering was used to group these peaks into three clusters (C1-C3) using concatenated data from men and women. Color bar on the top represents discrete age groupings as defined in this study (young, middle-aged, older). Rows are annotated according to their position relative to the nearest TSS: proximal if <1kbp distance, distal <u>otherwise</u>. (**b-c**) Annotations of shared temporal peaks using chromHMM states (**b**) and gene sets from DICE database^10^ (**c**). Colors represent hypergeometric enrichment test p-values; <u>light grey cells in plot indicate that there were insufficient genes in the gene sets to run an enrichment test.</u> The enrichment pattern strongly associates cluster C1 to T cells, suggesting a delayed loss of accessibility in women relative to men; C2 to CD19^+^ cells, suggesting the presence of CD19^+^ specific loci with opposing temporal behavior in men and women; and C3 to monocytes and NK cells. (**d**) Transcription factor (TF) motif enrichment results for each temporal cluster (C1-C3), relative to the other two clusters. Motif enrichment analyses carried out on 1,388 PWMs, grouped into families based on the sequence similarity, and most significant p-value for each motif family is represented here. <u>Tests were done using HOMER</u>. (**e-f**) Expression levels of TFs associated to cluster 1 (C1) and cluster 3 (C3) grouped by age group and sex <u>whose expression follows the same pattern as the peak temporal clusters where they are enriched</u>. Cluster 2 (C2) is omitted since <u>all TFs in this group show a significant increase with age in females or both sexes</u>. Box plots represent median <u>and IQR values, with whiskers extending to 1.5 times the IQR. Wilcoxon rank-sum test used to compare expression levels between sexes (significance value below boxes) and age groups (above boxes): *p<0.05, **p<0.01, ***p<0.001, ns: non-significant. Sample sizes for young individuals n=11F, 6M, middle-aged n=10F, 20M, older n=13F, 14M.</u> Source data are provided as a Source Data file for panels a,b,d.

These data revealed that the similarity of the aging signatures <u>between sexes</u> depends on the cell type. Age-related changes in naïve T cells (loss) and NK and memory CD8^+^ T cells (gain) were similar. However, gains in monocyte-specific loci were greater in men, which contributed to the sex-differences. B cell-specific loci lost accessibility/expression with age only in men, representing the most sex-dimorphic aging pattern.

### Chromatin accessibility and gene expression changes over adult lifespan

We uncovered chronological trends in epigenomic/transcriptomics maps using Autoregressive Integrated Moving Average (ARIMA)^25^ models by first finding the best-fitting ARIMA model to each peak/gene as a function of age and selecting the ones that displayed a chronological trend significantly different than random fluctuations. This uncovered 13,297 and 13,295 ATAC-seq peaks with a temporal trend in women and men, respectively, grouped into three major clusters (**Figure 4a**, Supplementary **Table 9**). Cluster1 (F1/M1) included peaks losing accessibility with age, whereas cluster2 and cluster3 included peaks gaining accessibility with age (F2/M2 and F3/M3). In cluster3, the gains occurred earlier (∼10-20 years) than in cluster2.

Enrichment analyses revealed that, in both sexes, cluster1 was associated to T cell-related loci, particularly naïve T cells (**Figure 4b**, Supplementary **Figure 6a**). Interestingly, cluster1 was significantly enriched in B cell-specific regions only in men, reinforcing a male-specific decline in B cells. Cluster2 was most significantly enriched in monocyte-specific regions in both sexes, albeit more significantly in men. Finally, cluster3 was most significantly enriched in NK cell-specific regions in both sexes, more significantly in men. Although less significant, we also detected female-specific enrichment of B cell-specific regions in cluster2 and cluster3, contrasting the declines in men (**Figures 2c, 4b**).

**Figure 6.**
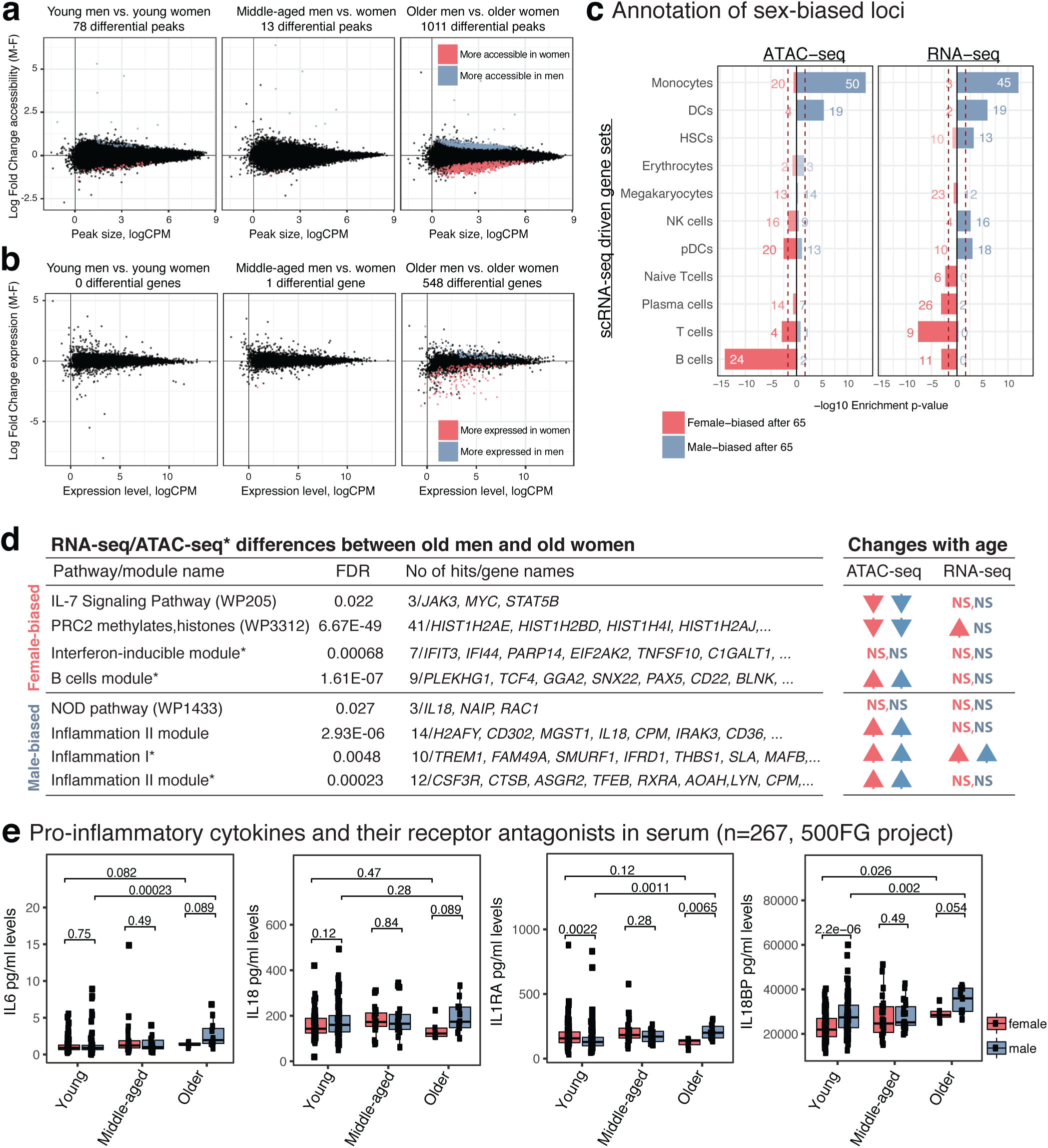
<u>Genomic differences between sexes at different age groups</u>. MA plots representing mean log2 fold change (male-female) versus average <u>log2 read counts per million reads (logCPM)</u> for ATAC-seq (**<u>a</u>**<u>) and RNA-seq (</u>**<u>b</u>**) data at three age groups using <u>78,167</u> peaks <u>(12,199 genes) at</u> autosomal chromosomes. <u>Both epigenomic and transcriptomic</u> differences between sexes increase with age. Peaks <u>and genes significantly upregulated</u> in women (men) <u>relative to men (women)</u> are represented in red (blue). <u>Fold changes obtained via GLM modeling of the data sets; statistical significance assessed at a 5% FDR threshold based on Benjamini-Hochberg p-value adjustment. Tests based on n=100 (ATAC-seq) and n=74 (RNA-seq) independent samples.</u> (**c**) Enrichment of <u>significantly</u> sex-biased peaks/genes in older individuals using cell-specific gene sets obtained from single-cell RNA-seq data. Note the male bias toward increased accessibility/expression for monocytes and DCs and the female bias toward increased accessibility/expression for T and B cells. P-values are based on hypergeometric enrichment tests. Numbers on bars represent the number of <u>differential</u> genes <u>overlapping</u> each gene set. (**d**) Left: Selected pathways/module enrichments for male-biased and female-biased genes/loci. The complete list of enrichments is presented in Tables S12, S13. * represents annotations obtained from ATAC-seq data analyses. Right: Arrows indicating whether the same pathway/module has been significantly associated with age-related changes. Significant enrichments (FDR 5%) and their directions are represented with arrows. Red arrows for women, blue arrows for men. NS = not significant. The complete list of enrichments is presented in Tables S6, S8. (**e**) ELISA data from 500 Functional Genomics (500FG) project to measure serum levels of pro-inflammatory cytokines (n=267). Protein expression levels are compared between sexes and between young and older individuals. P-values were calculated using <u>Wilcoxon rank sum</u> (two sided). Box plots represent <u>median and IQR values, with whiskers extending to 1.5 times the IQR.</u> Source data are provided as a Source Data file for panels a,b,c.

ARIMA models revealed temporal transcriptomic patterns consistent with the epigenomic patterns. 1,068 and 1,471 genes had temporal trends across human lifespan respectively in women and men (**Figure 4c**, Supplementary **Table 9**). We grouped these genes into three clusters (men: M1-3, women F1-3) based on the similarity of trends. Enrichment analyses^10^ revealed changes shared between sexes including down-regulation of naïve T cell genes and up-regulation of monocyte, NK, and effector T cell genes. Similar to the epigenomic trends, temporal genes of men and women differed for the B cell compartment, due to a male-specific downregulation of B and plasma cell-specific genes (**Figure 4d**, Supplementary **Figure 6b**) and a female-specific upregulation –albeit weaker– of plasma B cell-specific genes. These data suggest that, other than for B cells, temporal epigenomic/transcriptomic trends are shared between sexes and composed of genomic declines in T cell associated loci and increases in monocytes and NK cell associated loci.

### PBMCs go through rapid epigenomic changes at two discrete periods during adult lifespan

To study whether temporal epigenomic changes are acquired gradually over lifespan or more rapidly at certain ages, we detected age brackets during which abrupt changes take place, referred to as breakpoints. Within each temporal cluster (**Figure 4a**), we compared epigenomic profiles observed at ages immediately preceding and succeeding a given age (e.g., 5 years before/after) and calculated a p-value for differences between the windows. By using a moving window approach with varying window sizes, we ensured robustness to outliers, sample sparsity at certain ages, and temporal fluctuations (Supplementary **Figure 6c**). Finally, for each age, we calculated a combined p-value from different windows, where a smaller p-value indicates rapid epigenomic changes at that age. Due to RNA-seq sample sparsity, these analyses were restricted to ATAC-seq samples.

These analyses revealed two periods in adult lifespan during which rapid changes occur: i) a timepoint in late-thirties/early-forties, and ii) a later timepoint after 65 years of age (**Figure 4e**, Supplementary **Figure 6c**). The earlier breakpoint was detected between ages 38-41 in women across three clusters and ages 40-42 in men in cluster1 and cluster3. Magnitude (breakpoint p-values, **Figure 4e**) and timing of changes for this breakpoint were similar between sexes. Together with **Figure 4a**, this timepoint matched the start of aging-related epigenomic changes, i.e., the start of chromatin accessibility losses in cluster1 and gains in cluster2/3. The later breakpoint was detected between ages 66-71 in women across three clusters (**Figure 4e**, Supplementary **Figure 6c**). Interestingly, in men, this breakpoint was observed earlier (ages 62-64). The magnitude of changes for the second breakpoint was more pronounced in men than in women. Combined with **Figure 4a**, this breakpoint coincided with the acceleration of epigenomic changes, such as a faster loss of chromatin accessibility in cluster1. In summary, breakpoint analyses revealed that although aging-related changes accumulate gradually, there are at least two periods in the adult lifespan during which the immune system undergoes more abrupt epigenomic changes. The first breakpoint manifests itself in both sexes similarly, whereas the second latter breakpoint affects men more strongly and earlier than women. Timing of breakpoints were comparable across clusters, suggesting that cellular interactions might modulate these temporal trends.

### Shared temporal genomic patterns between men and women

We detected 3,197 peaks and 180 genes with shared temporal trends between sexes, which grouped into three clusters (**Figure 5a**, Supplementary **Figure7a**). Cluster1 captured a downward trend in both datasets and was associated with naïve T cells (**Figure 5b**, Supplementary **Figure 7b**). Similarly, cluster3 exhibited an upward trend and was associated to monocytes and NK cells. Interestingly, cluster 2 exhibited opposite temporal patterns between sexes: a downward trend in men and an upward trend in women and was associated to B cells (**Figures 5b, 5c**, Supplementary **Figure 7b**).

Further inspection of clusters identified temporal changes for important molecules. Cluster1 included T cell development and signaling genes, e.g., co-stimulatory molecule *CD27* and *TCF7* - a key TF in T cell development (Supplementary **Table 9**). Expression of both molecules decreased with age, the decrease was faster in men (Supplementary **Figure 7d**). In contrast, cluster2 exhibited contrasting temporal patterns between sexes and was associated to B and plasma cells, in both ATAC-seq and RNA-seq (**Figure 5b**, Supplementary **Figure 7b**). These included the E-protein *TCF4* (E2.2) that control various aspects of B cell development^26^ and *ORAI2* in the B cell receptor-signaling pathway; expression levels of both genes increased in women and decreased in men. Changes in cluster3 included up-regulation of *JAG2* in both sexes, which is expressed specifically in NK cells^10^ and plays critical roles in DC-mediated NK cell cytotoxicity^27^ (Supplementary **Figure 7d**).

To uncover potential regulators of epigenomic changes, we identified TF motifs enriched in temporal peaks using 1,388 motifs for 681 TFs grouped into 278 families. 50 of these families were significantly enriched across three clusters (**Figure 5a)**. These included families that were enriched across all clusters (e.g. a bHLH family motifs for TCF3, TCF4, TAL1) and enriched in a single cluster (**Figure 5d**, Supplementary **Table 10**, see Supplementary **Figure 7e** for sex-specific analyses). TFs associated to naïve T cells^10^ were enriched in cluster1 (e.g., NPAS2, TCF7, SOX12, FOXO1). Among these, TCF7 is a potential ‘pioneer factor’ and can impact the chromatin accessibility levels^28^. Expression levels of these TFs also decreased with age (**Figure 5e**). Cluster3 was enriched for motifs for MAFB, CEBPA, CREB5, and NFE2, which are highly expressed in monocytes^10^ and were upregulated with age in PBMCs (**Figure 5f**). TF enrichments for cluster2 were not as significant due to the size of this cluster. Combined analyses of temporal trends in men/women uncovered shared changes: i) downward genomic trends in naïve T cell genes/TFs, including the pioneer factor TCF7; ii) upward trends for genes/TFs active in monocytes and cytotoxic cells. These analyses further highlighted the stark differences between sexes regarding the aging of the B cell compartment.

### PBMCs of men and women diverge with age

We compared ATAC-seq and RNA-seq maps of men and women to describe sex-bias in human PBMCs at different age groups. In young and middle-aged subjects, very few differences were detected between sexes. Interestingly, differences between sexes increased after the age of 65, where 1011 differential peaks and 548 differential genes were detected (**Figures 6a, 6b**). Functional annotations of sex-differences were conserved between the two assays, revealing a bias for myeloid lineage particularly for monocytes in older men and a bias towards adaptive cells (B, T) in older women (**Figure 6c**, Supplementary **Figure 8a**, Supplementary **Table 11**). Epigenomic enrichments^20^ (Supplementary **Figures 8b, 8c**) confirmed that male-biased peaks were enriched in monocyte-specific loci and female-biased peaks were enriched in B/T cell-specific loci, including important signaling molecules (*JAK3, STAT5B*) (Supplementary **Table 11**). Furthermore, IL-7 signaling and other B/T cell signaling pathways were more active in PBMCs of older women compared to older men (Supplementary **Figure 6d**, Supplementary **Tables 12, 13**). Given that this pathway is downregulated with age in both sexes (**Figure 6d**, right table), the difference between older subjects suggest that the age-related decline in T cells is greater in men. In contrast, older men had higher activity for inflammation-related modules compared to older women, including the pro-inflammatory cytokine *IL18*. Chromatin accessibility levels for inflammation related genes/loci increases with age in both sexes (**Figure 6d**, right table), suggesting that although inflammation increases with age in both sexes, the magnitude is greater in men. In agreement, serum protein measurements from the 500 Human Functional Genomics consortium^11^ (n=267) revealed that older men have higher levels of pro-inflammatory proteins (IL18, IL6) <u>and their receptor antagonists</u> (IL18BP, IL1RA) compared to older women. IL18BP and IL6 levels increased with age in both sexes, however, the increase was greater in men leading to differences between older adults. In contrast, serum IL1RA levels increased specifically in men, also leading to a significant difference (<u>Wilcoxon</u> p=0.0065) between older men and women (**Figure 6e**).

These data suggest that older men show a bias towards innate immunity whereas older women towards adaptive immunity. This partially stems from the elevated activation of monocyte related genes/loci with age in men (**Figure 2c**, Supplementary **Figure 4c**). Fifteen times more monocyte-specific loci were activated in men with age (960 vs. 64 peaks). (Supplementary **Figure** 2f). However, we did not detect significant sex differences in CD14^+^ and CD16^+^ monocyte cell numbers (counts and frequencies) in flow cytometry data from Milieu Intérieur Consortium^12^ (n=892) (Supplementary **Figure 8d**) or from our cohort (**Figure 1d**). Even though the number of CD16^+^ monocytes increase with age in both sexes (Supplementary **Figure 8e**), this increase was similar between sexes (Supplementary **Figures 8d, 8e**). In contrast, the female-bias in adaptive immunity partially stemmed from male-specific genomic declines in B cell associated loci (**Figures 2c, 4d**) and B cell percentages within PBMCs (**Figure 1d**). This trend was also observed in the Milieu Intérieur cohort for different B cell subsets (i.e., naïve, transitional, founder) (Supplementary **Figures 8d, 8e**). Interestingly, mass-cytometry data from a third cohort^29^ (n=190 Caucasians) also revealed significant age-related declines in B cell percentages (Supplementary **Figure 8f**). However, these declines were observed in both sexes. Further studies are needed to uncover whether B cell related differences among cohorts can be due to clinical and/or environmental factors.

## Discussion

Despite well-characterized sex differences in immune responses, disease susceptibility, and lifespan, it is unclear whether aging differentially affects peripheral blood cells of men and women. To fill this gap, we generated ATAC-seq, RNA-seq, and flow cytometry data in PBMCs from 172 age-matched healthy adults. Using novel systems immunology pipelines, we discovered a genomic signature of aging that is shared between sexes including 1) declines in T cell functions, 2) increases in cytotoxic (NK, memory CD8^+^ T) and monocyte cell functions. This signature is in alignment with known age-related changes in the immune system including declining adaptive responses and increased systemic inflammation with age^14, 30^. Despite these similarities, male PBMCs went through greater changes that are not attributable to clinical differences. Notably, 15 times more monocyte-specific loci were activated in men compared to women. Previous studies in whole blood showed that DNA methylomes age faster in men compared to women^31^. Here, we uncovered an accelerated aging phenotype for men in ATAC-seq data and uncovered cell types, cellular functions, and molecules that differentially age between sexes. In addition, our data revealed for the first time that aging has opposing effects on the B cells of men and women, where B cell-specific loci/genes were modestly activated in women but significantly inactivated in men, which might contribute to sex-differences in auto-immunity and humoral responses.

We studied middle-aged individuals to capture chronological changes and the timing of changes, which is essential in distinguishing age-related changes from those resulting from maturation^32^. Chronological patterns in ATAC-seq and RNA-seq data were correlated and included increasing accessibility/expression for cytotoxic cells and monocytes, and decreasing accessibility/expression for T cells in both sexes. Temporal patterns from men and women further highlighted the differential aging of the B cell-related loci. Breakpoint analyses uncovered that although aging-related epigenomic changes accumulate gradually throughout adult life, there are two periods in the human lifespan during which the immune system undergoes abrupt changes. The first breakpoint was similar between sexes, however the second breakpoint was detected earlier in men (5-6 years), which is comparable to the life-span differences: 76.9 for men and 81.6 for women in USA^33^. In both sexes, the second breakpoint was associated with accelerated epigenomic changes and occurred ∼12-15 years before the end of the average lifespan. The differences in the timing of age-related changes can be helpful in clinical decisions regarding when to start interventions/therapies.

PBMCs of men and women significantly differed after the age of 65, contrary to our expectations due to declining sex hormones. Annotation of sex-biased loci revealed that older women have higher genomic activity for adaptive cells and older men have higher activity for monocytes and inflammation. Some of these differences, but not all, are attributable to changes in cell compositions, e.g., the decline of B cell frequencies in older men (**Figure 1d**), which is confirmed in a second cohort^12^ (Supplementary **Figure 8d**) and has been reported in a Japanese cohort^34^. This concordance across cohorts suggests that sex-specific aging signatures described here might be conserved across ethnicities/populations, which requires further investigation. Age-related increases were associated with inflammatory pathways/genes more significantly in men, suggesting an accelerated inflamm-aging signature for men. Data from an independent cohort^11^ supported this, where older men had higher levels of pro-inflammatory cytokines (IL6, IL18) in their serum compared to older women (**Figure 6e**). Interestingly, we did not detect significant differences between older men and older women in CD14^+^ and CD16^+^ monocyte cell numbers/frequencies neither in this cohort nor in a separate bigger cohort^12^ (Supplementary **Figure 8e**). Therefore, age-related activation and sex-differences in monocytes potentially stems from cell-intrinsic changes. Future studies are needed to explore reasons behind sex differences in inflammaging, including the role of sex hormones and differential activation of DAMPs/PAMPs or transposable elements^35^.

Age-related changes in DNA methylation levels are predictive of ages across tissues and conditions, most notably the Horvarth’s clock including 353 CpG sites^36^. A total of 187 (53%) of these CpG sites overlapped PBMC ATAC-seq peaks (86,145). The missing overlap likely stems from differences in technologies: CpG arrays versus sequencing-based ATAC-seq assay that genome-wide capture active regulatory elements^37^. Only a few CpGs in the clock overlapped opening/closing peaks (Supplementary **Table 16**), suggesting that ATAC-seq and DNA methylation arrays capture distinct epigenomic information and aging signatures. ATAC-seq captures age-related changes in regulatory element activity in a cell-specific manner. Although the mechanism underlying the CpG clocks are not established with certainty, they are proposed to represent widespread entropic decay of the DNA methylation landscape^38^.

Using a systems immunology approach in human PBMCs, this study uncovered cell types and immune functions that are differentially affected with aging between men and women. Changes in bulk PBMCs were annotated using cell-specific regulatory element loci inferred from reference epigenomic datasets^10,20^. Although this approach was effective in annotating the aging signatures, it is prone to biases in references, e.g., differences in data quality and limitation to use cell types available in references. Future studies are needed to describe these sex differences at single-cell resolution and in sorted cells and to establish their functional implications. Moreover, future studies are needed to study important molecules identified here (*IL7R, LEF1, TCF7, IL8, IL18*) as sex-specific biomarkers of immune system aging. Taken together, these findings indicate that sex plays a critical role in human immune system aging and should be taken into consideration while searching for molecular targets and time frames for interventions/therapies to target aging and age-related diseases.

## Materials and Methods

### Human Subjects

All studies were conducted following approval by the Institutional Review Board of UConn Health Center (IRB Number: 14-194J-3). Following informed consent, blood samples were obtained from 172 healthy volunteers residing in the Greater Hartford, CT, USA region recruited by the UConn Center on Aging Recruitment and Community Outreach Research Core (http://health.uconn.edu/aging/research/research-cores/). For older adults 65 years and older, recruitment criteria were selected to identify individuals who are experiencing “usual healthy” aging and are thus representative of the average or typical normal health status of the local population within the corresponding age groups^5^ Selecting this type of cohort is in keeping with the 2019 NIH Policy on Inclusion Across the Lifespan (NOT-98-024)^39^, increasing the generalizability of our studies and the likelihood that these findings can be translated to the general population^40^. Subjects were carefully screened in order to exclude potentially confounding diseases and medications, as well as frailty. Individuals who reported chronic or recent (i.e., within two weeks) infections were also excluded. Subjects were deemed ineligible if they reported a history of diseases such as congestive heart failure, ischemic heart disease, myocarditis, congenital abnormalities, Paget’s disease, kidney disease, diabetes requiring insulin, chronic obstructive lung disease, emphysema, and asthma. Subjects were also excluded if undergoing active cancer treatment, prednisone above 10 mg day, other immunosuppressive drugs, any medications for rheumatoid arthritis other than NSAIDs or if they had received antibiotics in the previous 6 months. Beyond these steps to exclude specific chronic conditions we also undertook further additional efforts to exclude older adults with any significant frailty. Since declines in self-reported physical performance are highly predictive of frailty, subsequent disability and mortality^41^, all subjects were also questioned as to their ability to walk ¼ mile (or 2-3 city blocks). For those who self-reported an inability to walk ¼ mile^41^, the “Timed Up and Go” (TUG) test was performed and measured as the time taken to stand up from the sitting position, walk 10 feet and return to sitting in the chair^42^. Scoring TUG > 10 second was considered an indication of increased frailty and resulted in exclusion from the study^43^. Information on medications in **Table S1** illustrates that as expected medication usage did increase with age. Nevertheless, these medications all reflected their use for common and controlled chronic conditions unlikely to influence our findings such as hypertension, hyperlipidemia, hypothyroidism, degenerative joint disease, seasonal allergies, headaches, atrial fibrillation, depression, anxiety, or ADHD (attention deficit hyperactivity disorder). Finally, smoking history data are not typically collected in these studies –including ours- since smoking is a rare habit among older adults.

### Ethics

The study was conducted following approval by the Institutional Review Board of UConn Health Center (IRB Number: 14-194J-3). All study participants provided written informed consent at baseline using institutional review board-approved forms. <u>Individual-level human genomic data (ATAC-seq and RNA-seq) are shared in a federal controlled-access database through dbGaP.</u> Our study complies with Tier 1 characteristics for “Biospecimen reporting for improved study quality” (BRISQ) guidelines.

### Flow cytometry data generation and analyses

PBMCs were isolated from fresh whole blood using Ficoll-Paque Plus (GE) density gradient centrifugation. For the analysis of the frequencies of Naïve T cells (CD45RA^+^CCR7^+^), Central Memory T cells (CM; CD45RA^-^CCR7^+^), Effector Memory T cells (EM; CD45RA^-^CCR7^-^), and Effector Memory RA (EMRA; CD45RA^+^CCR7^-^), B cells and Monocytes, PBMCs were stained with fluorochrome-labeled antibodies specific for <u>CD3 (UCHT1) Biolegend-1:100 (cat# 300436), CD4 (RPA-T4) Biolegend 1:80 (cat# 558116), CD8 (SCFI21Thy2D3) Beckman Coulter 1:80 (cat# 6604728), CD45RA (HI100) BD biosciences 1:80 (cat# 560674), CD19 (HIB19) BD biosciences 1:100 (cat#-555415), CD14 (MSE2) BD biosciences 1:80 (cat# 557923), CCR7 (150503) BD biosciences 1:20 (cat# 561271).</u> The stained cells were acquired with BD Fortessa and analyzed with FlowJo software (TreeStar). Flow data is analyzed using Generalized Linear models (GLM) to quantify the association between cell proportions and age, sex, and their interaction (age and sex). For the analyses of major cell populations, we used age (continuous variable) as a covariate, whereas for T cell subsets we used age group (old vs. young) as a variable. <u>We excluded one outlier individual (subject 104) from downstream analyses.</u>

### ATAC-seq library generation and processing

ATAC-seq^44^ data was generated from fifty thousand unfixed nuclei using Tn5 transposase (Illumina, Nextera DNA sample prep kit) for 30 min at 37°C. The resulting library fragments were purified using Qiagen MinElute kit (Qiagen). Libraries were amplified by 10-12 PCR cycles, purified using a Qiagen PCR cleanup kit (Qiagen), and finally sequenced on an Illumina HiSeq 2500 with a minimum read length of 75 bp to a minimum depth of 30 million reads per sample. At least two technical replicates (average = 2.4 replicates) were processed per biological sample. **Table S2** summarizes the depth, peak number, and fragments in reads (FrIP) scores for ATAC-seq samples. ATAC-seq sequences were quality-filtered using trimmomatic^45^, and trimmed reads were mapped to the GRCh37 (hg19) human reference sequence using bwa-mem^46^. After alignment, technical replicates were merged and all further analyses were carried out on these merged data. For peak calling, MACS2^47^ was used with no-model, 100bp shift, 200bp extension, and bampe option. Only peaks called with a peak score (q-value) of 1% or better were kept from each sample, and the selected peaks were merged into a consensus peak set using Bedtools multiinter tool^48^. Only peaks called on autosomal chromosomes were used in this study. We further filtered consensus peaks to avoid likely false positives by only including those peaks overlapping more than 20 short reads in at least one sample, and peaks for which the maximum read count did not exceed 500 counts per million (cpm) to account for regions that are potential artifacts. Finally, we excluded peaks overlapping ENCODE blacklisted regions downloaded from http://hgdownload.cse.ucsc.edu/goldenpath/hg19/encodeDCC/wgEncodeMapability/. An additional quality-control step was developed to filter out samples with a consistently poor signal, consisting of an algorithm to discover and characterize a series of relatively invariant benchmark peaks, defined as a set of peaks expected to be called in all samples. Samples that consistently miss calls for a significant portion of these benchmark peaks are flagged as having poor quality. A benchmark peak is defined based on three criteria, namely (1) that it remains approximately invariant between the two groups of interest (i.e., young and old samples), (2) that it captures a substantial number of reads, and (3) that it is called in most samples. For each peak, the absolute value of the log of the ratio of healthy old to healthy young mean normalized read counts (log fold change, logFC) was used to assess the first criteria, whereas the maximum read count over all samples (maxCt) is used to assess the second one. In this study, a peak was considered apt for benchmarking when (1) its absolute logFC was in the bottom quartile of the distribution over all peaks, (2) its maxCt was in the top decile of the distribution over all peaks, and (3) the peak was called in at least 90% of the samples. Using these parameters, 640 (out of 86,300) peaks were selected as benchmark; only samples for which at least 92.5% of these peaks were called were selected for analyses, which excluded 8 samples from further analyses. We examined the effects of each of these parameter choices and found that the same samples were consistently chosen as poor quality for a range of values chosen to assess the benchmark criteria. After re-applying the peak selection criteria to the remaining 100 samples, we arrived at a peak count of 86,145 peaks. Among the samples that passed the QC step, we studied the distribution of FRIP scores, depth of sequencing, and peak numbers (that are correlated measures), which revealed a wide range of values. Neither of these measures were correlated significantly with age in linear regression models (FRIP: <u>Pearson</u> *<u>r</u>* = 0.11, p = 0.29; Depth: <u>Pearson</u> *<u>r</u>* = 0.081, p = 0.42; Peak number: <u>Pearson</u> *<u>r</u>* = 0.1, p = 0.32). However, on the average samples from men had higher values compared to samples from women (FRIP: 0.218 vs. 0.178; Depth: 103M vs. 72M; Peak number: 37046 vs. 29170). To account for these differences, prior to statistical analyses, ATAC-seq read counts were normalized to each sample’s effective library size (i.e., the sum of reads overlapping peaks) using the trimmed mean of M-values normalization method (TMM)^49^. In addition, we used the effective library size and significant surrogate variables as co-variates in differential analyses (see Differential analyses for details). Accordingly, we noted that SV1 in these analyses correlate significantly with library depth (<u>Pearson</u> *r*=-0.68 p=6.8e-15) and number of peaks (<u>Pearson correlation</u> *r*=-0.31, p=0.0016) in old-young comparisons and with library depth (<u>Pearson</u> *<u>r</u>*=-0.56, p=1.7e-09) and FRIP score (<u>Pearson</u> *<u>r</u>*=0.2, p=0.044) in men-women comparisons.

### RNA-seq library generation and processing

Total RNA was isolated from PBMCs using the Qiagen RNeasy (Qiagen) or Arcturus PicoPure (Life Technologies) kits following manufacturer’s protocols. During RNA isolation, DNase I treatment was additionally performed using the RNase-free DNase set (Qiagen). RNA quality was checked using an Agilent 2100 Bioanalyzer instrument, together with the 2100 Expert software and Bioanalyzer RNA 6000 pico assay (Agilent Technologies). RNA quality was reported as a score from 1 to 10, samples falling below threshold of 8.0 being omitted from the study. cDNA libraries were prepared using either the TruSeq Stranded Total RNA LT Sample Prep Kit with Ribo-Zero Gold (Illumina) or KAPA Stranded mRNA-Seq Library Prep kit (KAPA Biosytems) according to the manufacturer’s instructions using 100ng or 500ng of total RNA. Final libraries were analyzed on a Bioanalyzer DNA 1000 chip (Agilent Technologies). Paired-end sequencing (2×100bp) of stranded total RNA libraries was carried out in either Illumina NextSeq500 using v2 sequencing reagents or the HiSeq2500 using SBS v3 sequencing reagents. Quality control (QC) of the raw sequencing data was performed using the FASTQC tool, which computes read quality using summary of per-base quality defined using the probability of an incorrect base call^50^. According to our quality criteria, reads with more than 30% of their nucleotides with a Phred score under 30 are removed, whereas samples with more than 20% of such low-quality reads are dropped from analyses. Benchmarking is also applied on RNA-seq data using the same benchmark parameters as ATAC-seq, which resulted in 304 benchmark genes, none of the RNA-seq samples were dropped due to poor quality. Reads from samples that pass the quality criteria were quality-trimmed and filtered using trimmomatic^45^. High-quality reads were then used to estimate transcript abundance using RSEM^51^. Finally, to minimize the interference of non-messenger RNA in our data, estimate read counts were re-normalized to include only protein-coding genes. Table S2 summarizes the quality control measures for our PBMC RNA-seq samples.

### Differential analysis

To identify differentially open chromatin regions from ATAC-seq and differentially expressed genes from RNA-seq data, the R package edgeR was used to fit a generalized linear model (GLM) to test for the effect of aging between healthy young and healthy old samples by sex, as well as the effect of sex by age group. In addition to sex and age group (old vs. young), our models included the base-2 log of effective library size to ensure peakwise normalization. We isolated a batch effect correlated to time period whereby samples were collected and libraries were prepared, and used ComBat to adjust the data for this effect. Finally, we used Surrogate Variable Analysis (SVA^52^) to capture unknown sources of variation (e.g., localized batch effects, subject-level heterogeneity, variation in library preparation techniques) statistically independent from age group assignments. SVA decomposes the variation that is not accounted for by known factors like age group or sex, into orthogonal vectors that can then be used as additional covariates when fitting a model to test for differential accessibility or expression. Using the built-in permutation-based procedure in the R package sva, we choose to retain one SV to include as covariate in the GLM model for PBMC ATAC-seq and none for RNA-seq data analyses^53^. GLM models where implemented using a negative binomial link function, including both genome-wide and peak-specific dispersion parameters, estimated using edgeR’s “common,” “trended,” and “tagwise” dispersion components, calculated using a robust estimation option. Benjamini-Hochberg P-value correction was used to select differentially open peaks at a False Discovery Rate (FDR) of 5%. To generate a set of model-adjusted peak estimates of chromatin accessibility (i.e., batch-, and SV-adjusted) for downstream analyses and visualization, we used edgeR to fit a “null” model excluding the sex and age group factor, and then subtracted the resulting fitted values from this model from the original TMM-normalized reads.

### Peak annotation and downstream analyses

Multiple data sources were used to annotate ATAC-seq peaks with regard to functional and positional information. HOMER^54^ was used to annotate peaks as “promoter” (i.e., within 2 kb of known TSS), “intergenic”, “intronic”, and other positional categories. For functional annotation of peaks, we used a simplified scheme integrating public chromatin states calculated for major PBMC subpopulations with ChromHMM from Roadmap Epigenomics^20^, Blueprint Epigenome, and a third reference data^55^: First, we intersected the chromHMM-generated states with our set of consensus peaks, and solved conflicting cases where multiple chromatin states overlap the same ATAC-seq peak so that each peak was assigned a single annotation, according to the following priority rules: if a peak overlaps both an active TSS and enhancer region, which state takes priority depends on whether the peak is proximal (i.e., within 1,000 bp of the nearest TSS), in which case it is annotated as a promoter, or distal (distance to nearest TSS greater than 1,000 bp), when it is annotated as an enhancer instead. For all other states, rules apply as follows: Active Enhancer > Genic Enhancer > Bivalent TSS > Weak Enhancer > Bivalent Enhancer > PolyComb repressed > TSS Flanking > Transcription > ZNF Genes and repeats > Heterochromatin > Quiescent/Low signal. Then, to facilitate interpretation and visualization, we simplified the original sets of chromatin states to a scheme with 6 pooled meta-states, namely (1) TSS, collecting active, flanking, and bivalent TSS states; (2) Enhancer, pooling active, weak and bivalent enhancer states; (3) Repressed PolyComb, combining both weak and strong PolyComb states; (4) Transcription, including both weak and strong transcription states, (5) the quiescent chromHMM state; and (6) other states (ZNF, heterochromatin) combined together. To annotate peaks as cell-specific for a given subset obtained from one of the three datasets listed above, we determined for each peak whether it was annotated as an active promoter or an active enhancer in a single cell population or lineage, and in such cases labeled the peak accordingly as cell- or lineage specific. For example, if a peak is annotated as an active enhancer in both naïve and memory CD4^+^ T cells but as another state (e.g., repressed) in every other subset, then the peak would be considered CD4^+^ T-specific. For gene-based analyses, HOMER was used to assign each ATAC-seq peak to the nearest TSS, as measured from the peak center. To improve confidence on these annotations, gene-based analyses were further restricted to include only peaks located within 100 kb of their corresponding TSS. ATAC-seq peaks were also annotated using gene sets provided by curated immune function-related co-expression modules^13^. These gene sets comprise 28 modules defined from multiple compiled transcriptomic data sets, which were originally annotated based on expert knowledge of representative functions of the gene ensemble in each module. In this study, we have preserved and used these annotations to test for enrichment of these modules in gene sets of interest, such as the set of genes associated to chromatin peaks gaining or losing accessibility with aging. We assessed enrichment using the hypergeometric test followed by Benjamini-Hochberg FDR adjustment for P-values. Further functional enrichment analyses were carried out using Wikipathways pathways^56^, immune modules^22^, gene sets from DICE database^10^ and single cell RNA-seq data in PBMCs. Gene sets from PBMC scRNA-seq data is driven from one-vs-all cell cluster comparisons. First, we identified 17 clusters from PBMC scRNA-seq data (n=26 samples) using the Louvain clustering in the ScanPy toolkit^57^. These clusters were manually inspected and assigned to different cell types by studying the expression of known marker genes. For each cluster, differentially expressed genes were identified using one-vs-all approach based on T-test followed by FDR. Marker genes for a cluster (or cell population) were found by using FDR <=5% and logFC > log2(1.25) using differential analyses results. Similarly, marker genes from the DICE database are obtained using their differential analyses results and the same cutoffs for cell-specific genes (FDR <=5% and logFC > log2(1.25)). Gene sets used in our differential analyses are provided as a supplementary table (**Table S14)**. Similarly, peak sets from ChromHMM states used in the enrichment analyses are provided as a supplementary table (**Table S15**). For each gene set, we tested for enrichment using the hypergeometric test, against a background defined by the set of genes that are expressed, as determined by RNA-seq data, or potentially expressed, as given by promoter accessibility, in PBMCs. We used the Benjamini-Hochberg FDR multiple test correction to assess significance of hypergeometric P-values.

### ATAC-seq and RNA-seq comparisons

Gene expression (RNA-seq, see above) data was generated for a subset of subjects with ATAC-seq profiles (summarized in **Table S1**). These data were normalized to protein-coding transcripts, and annotated to ENSEMBL GRCh37 gene symbols. Genes for which at least three normalized reads per million were obtained in at least two samples were considered as expressed, all others removed prior to analysis. This resulted in a total estimate of 14,157 expressed genes in PBMCs. We built a data set comprising paired ATAC-seq and RNA-seq samples by matching promoter peaks to nearest gene (TSS) annotations, defining promoters as the regions within 1000 bp flanks of each TSS. Whenever multiple expressed genes were matched to the same promoter peak, the peak with the maximum fold change was chosen for visualization.

### Inferring chronological aging trends

We used Autoregressive Integrated Moving Average (ARIMA) models as implemented in R on age-ordered normalized chromatin accessibility and expression data separately for each sex, to select genomic features (ATAC-seq peaks or mRNA-seq transcripts) to uncover significant chronological trends, as opposed to unstructured “noise” fluctuations undistinguishable from stationary data. Input data from all subjects in the study (see **Table S1**) were normalized as described above in “Differential Analyses”, i.e., corrected for known batch effects using ComBat, and model-adjusted to correct for library size and unknown batch effects (i.e., surrogate variables) using EdgeR. Thus, ARIMA-modeled data contains variation due to age, sex, and error variance unaccounted for by batch and library size effects. For model fitting, genomic features were converted into z-ordered objects (using R package zoo) after averaging values obtained for the same sex and chronological age (in years), and into time series objects. Formatted time series were then analyzed using auto.arima (from the forecast R package), using stepwise search and Akaike Information Criterion (AIC) to select the best model. This algorithm scans a wide range of possible ARIMA models and selects the one that satisfies the optimality criterion, i.e., the smallest AIC. Only non-seasonal models with a maximum difference (D) of 2 were considered in this study, such that a non-stationary trend the data can be modeled after integrating a stationary series once or twice. Overall, best-fitting ARIMA models based on chromatin accessibility or gene expression data included models with 0-7 AR, 0-5 MA, and 0-7 AR+MA parameters for accessibility and 0-4 AR, 0-3 MA, and 0-4 AR+MA parameters for expression. From these models, we chose for inspection only the ones carrying a “significant” trend, defined as those where (1) the difference order from stationary was higher than zero, and (2) at least one AR or MA coefficient was included in the model (i.e., D>0 ∩ AR+MA>0). These criteria allow sampling a wide variety of trend patterns if they were present in the data. For accessibility data, 14,594 (18.7%) and 13,591 (17.4%) peak models had D=1 (none had D=2) in females and males respectively, out of which 13,297 (17%) and 13,295 (17%), respectively, also were estimated to have AR+MA>0, and hence chosen for further analyses. Out of the chosen peaks, 12,941 and 11,566 had only MA (9,082 in females, 8,696 in males) or AR (2,562 in females, 2,574 in males) coefficients, whereas only 4,196 (31.6%) and 4,181 (31.4%) were estimated to have two or more non-zero (AR+MA) coefficients in females and males, respectively. A total of 3,197 (4.1%) peaks were chosen in both sexes, whereas the remaining peaks were treated as sex-specific. For expression data, 1,367 (11.1%) and 1,665 (13.5%) models had D=1 (none had D=2) in females and males respectively, out of which 1,068 (8.6%) and 1,471 (11.9%), respectively, also were estimated to have AR+MA>0, and hence used in further analyses. Out of these, 1,359 and 832 had only MA (511 in females, 903 in males) or AR (480 in females, 372 in males) coefficients, whereas only 503 (36.8%) and 554 (33.3%) were estimated to have two or more non-zero (AR+MA) coefficients in females and males, respectively. A total of 180 (1.5%) peaks were chosen in both sexes, whereas the remaining peaks were treated as sex-specific.

### Temporal peak/gene analyses

After selecting genomic features with a significant trend, we used their estimated parameters to fit the original data to reduce noise and facilitate the discovery of common trend patterns across multiple features. Fitted values were computed at the same chronological ages observed in the data for each sex, converted to *z*-scores for comparability, and then submitted to *k*-means clustering for aggregation into *k* sets of similar chronological aging patterns. Chosen features for each sex were clustered separately, and additionally features chosen as common to both sexes were concatenated as a single sequence of two age-ordered ages prior to normalization. This concatenation approach was designed to facilitate the detection of features with consistent chronological patterns across sexes. For clustering, we used the R-stats version of the *k*-means algorithm (kmeans), starting from nstart=1,000 random sets of *k* cluster centers and running each for max.iter=100 iterations. To choose *k*, we opted for a biologically informed criterion as opposed to a purely statistical one, after noting that criteria such as maximization of within- vs. between-cluster variance tended to select a large (>20) number of slightly different clusters, likely emphasizing small differences in ARIMA model parameters with little biological significance. Instead, we computed optimal clusters for values of *k* between 2 and 15 and calculated enrichment statistics for cell-specific features (i.e., cell-specific chromatin states for ATAC-seq, and cell-specific expressed genes for RNA-seq data) for each cluster relative to all trending features, and asked whether adding clusters to the *k*=2 cluster case resulted in a gain of information demonstrated by cluster enrichment patterns qualitatively different from the patterns observed in absence of the extra clusters. In some cases, finding a cluster configuration satisfying this condition of biological discrimination required applying hierarchical clustering to collapse distinct *k*-means clusters <u>with the same biological signal</u>. For chromatin accessibility, conservative application of these criteria led to the selection of *k*=5 and *k*=6 clusters for females and males, respectively, which were collapsed into <u>3</u> clusters in both cases (one closing, two opening with aging), and *k*=3 for peaks trending in both sexes (one closing and one opening in both sexes, plus one opening in females while closing in males). The same numbers of final clusters were defined based on gene expression.

### TF motif enrichment analyses

To further explore functional associations of these clusters, we tested for enrichment of known and de novo transcription factor (TF) motifs in each chromatin accessibility cluster for female, male, and sex-pooled peak data, relative to all trending clusters for the same set of peaks. We used <u>the software</u> HOMER^54^ to both detect de novo motifs and to test for enrichment of 1,388 motifs obtained from JASPAR CORE 2018 database^58^ and from Jolma et al^59^ SELEX-derived <u>position weight matrices</u> (PWMs), corresponding to 680 distinct TFs. We used HOMER to assign a TF to each de novo motif found, based on sequence similarity to the 1,388 known motifs. From the pooled set of known and de novo motifs tested for each cluster, we used Benjamini-Hochberg method on enrichment p-values to estimate false discovery rates (FDR), and applied a 5% FDR significance threshold to select for the final set of enriched motifs. Since some motifs have similar consensus sequences and PWMs, prior to visualization of these results we grouped motifs into “families” defined as sets of motifs with nearly identical sequences, as determined by pairwise comparison of all 1,388 motif using MEME/TOMTOM^60^, which tests whether two motif sequences are more different than expected by chance. Using stringent 0.001% q-value and 5 E-value cutoffs, we identified 429 motif families with extremely similar consensus sequences, 319 of which were singletons (maximum family size=226 motifs).

### Breakpoint analyses

We investigated systemic chronological signatures in temporal peaks by testing, in each cluster, for the existence of “breakpoints,” i.e., short age intervals characterized by significant differences in accessibility in the intervals preceding and following the age interval. For each age *t* in the sampled age interval from *t*_min_ to *t*_max_, we tested for mean difference in accessibility between subjects with ages in the intervals *t*_min_-*w* vs. *t*_min_+*w*, where *w* represents a variable window span parameter, and plotted the observed p-values (i.e., -log10 P, or *loginvp*) as a function of age (*t*) to identify maxima that suggested the presence of discrete breakpoints. These tests were carried out on normalized and model-adjusted accessibility data corresponding to the ATAC-seq peaks associated to each sex-specific and common cluster identified as trending by ARIMA as described above. Since there were many more peaks than subjects for any given comparison, we used PCA to reduce the dimensionality of each cluster to n=3 PCs, and used MANOVA on these three dimensions to compute p-values at each tested value of *t*. For any given value of *w*, offset values for *t*_min_ and *t*_max_ were adjusted to match the age of available samples in the study. For example, a window span of 5 years required a *t*_min_ =27 if the youngest available subject was 22 years old, and a *t*_max_ =83 if the oldest available subject was 90 years old. For a given value of *w*, results from tests contrasting younger vs. older intervals would vary depending on sample size volume and imbalance, with statistical power increasing with the size of the window span. To take advantage of this effect, we deployed a multi-scale algorithm where we carried out tests using *w* values ranging from 10 to 20 years in order to identify breakpoints that were maximally supported under multiple window spans. Due to sample sparsity and variation, however, tests carried out under varying values of *w* may be unevenly affected by edge effects and influencing outlier points, which may result in strong significance of a comparison because of the presence of outliers and the partial overlap of a sampled interval with a breakpoint, limiting the ability of the method to precisely discover where such breakpoints may lie. To limit these effects and increase the robustness of the tests, we smoothed the *loginvp* distributions by fitting LOESS regressions to each comparison (i.e., each set of tests with the same *w* value) under a range of smoothing bandwidth parameters (i.e., bw = 0.25, 0.30, …, 0.70, 0.75). We combined the resulting eleven p-values at every sampled age using the Fisher’s method, reapplied LOESS smoothing to the resulting distribution, and used numerical differentiation to determine whether each age was predicted to be minimum or a maximum. Finally, we marked every maximum as a significant breakpoint candidate if it satisfied both a parametric criterion, i.e. significance of the Fisher method-combined p-values (χ^2^ test), and a heuristic criterion, namely whether the distance between this local maximum and the nearest minimum equaled or exceeded 25% of the value of the global maximum. The procedure described above results in a smoothed *loginvp* distribution for each *w* value, each comprising a series of points including maxima and minima, such that slightly different maxima can be estimated for different w values. Finally, we used Gaussian mixture modeling on the distribution of these maxima, as implemented in R-Mclust package, to group *loginvp* maxima obtained from different window spans into cohesive breakpoint intervals, whose medians and ranges we report herein for each cluster. Since breakpoints are independently calculated for each cluster, observed overlaps are likely the result of aging-related events with a genome-wide impact.

### Analyses of 500FG and MI data

We obtained publicly available ELISA data measuring serum protein levels by the 500 Human Functional Genomics (500FG) consortium^11^. We only retained individuals who are matching our cohort in terms of the age span, which resulted in data from 267 individuals. These individuals are grouped together using the age brackets defined in our study: Young 22-40 years old, middle-aged: 41-64 years old, older: 65 years old. We compared data from 1) men and women at all age brackets; 2) young men to old men; and 3) young women to old women using Wilcoxon Rank Sum non-parametric test (two sided). Note that flow cytometry data from this same cohort was not publicly available; hence cannot be included into the analyses. Similarly, publicly available flow cytometry data from Milieu Intérieur Consortium^12^ cohort was obtained. We used data that is already processed by this study and just used individuals whose ages are matching our cohort, which resulted in data from 892 individuals. We built linear models using R (lm function) to quantify the association between each flow cytometry measurement to age group (young, middle-aged, older), sex (F, M) and their interaction (age*sex). Significant associations with age (at FDR 5%) and with sex (at FDR 10%) are plotted in Figures S7D-E. No significant associations were detected with the interaction of sex and age at FDR 10%.

### R Shiny Application

The R Shiny package^61^ was used to create a webpage for the interactive visualization of the data from this study. Plots were generated using ggplot2^62^, using graph aesthetics used throughout the manuscript figures. Statistics for box plots were calculated using the wilcox.test function in R and presented without correction for the multiple (n = 5) comparisons (two sided). Statistics for scatterplots were calculated by fitting a linear model with the lm function in R using the formula (*measurement* ∼ *age*:*sex* + *sex*). For ATAC-seq data, only the peak closest to the gene TSS was considered.

## Author contributions

D.U., G.A.K. and J.B. designed the research. G.A.K coordinated the clinical sample collection. C.C., R.M., R.R., performed the experiments. E.M pre-processed the sequencing data. E.M, D.U., D.J.M., D.N., and A.E. analyzed the data. E.M., D.U. wrote the paper. D.J.M. implemented the R Shiny application. All authors revised the manuscript and helped with data interpretation.

## Acknowledgements

We thank Taneli Helenius for aid in scientific writing, research staff in the UConn Center on Aging for their help in recruitment and sample collection, JAX genomic technologies for their help with generating the sequencing data, and JAX Research IT for the support with building and maintaining the R Shiny application <u>as well as with data upload to dbGAP</u>. We thank members of Ucar, Stitzel, Bachereau labs, Michael Stitzel, Karolina Palucka, Virginia Pascual, Derya Unutmaz for critical feedback during the progress of the study. <u>We thank Yuqi Zhao and Magalie Collet for their help with dbGAP data upload.</u> This study was made possible by generous financial support of the National Institute of General Medical Sciences (NIGMS) under award number GM124922 (to DU) and National Institutes of Health (NIH) grants R01 AG048023, P01 AG021600, U19AI089987 (to JB). Opinions, interpretations, conclusions, and recommendations are solely the responsibility of the authors and do not necessarily represent the official views of the National Institutes of Health (NIH). G.A.K is supported by the Travelers Chair in Geriatrics and Gerontology.

## Data availability

The data generated as part of this study is controlled access. A subset of the ATAC-seq and RNA-seq samples used in these analyses was made public through EGA (Id: EGAS00001002605 [https://www.ebi.ac.uk/ega/studies/EGAS00001002605]). All PBMC ATAC-seq and RNA-seq samples used in this study can be found at dbGAP (Id: phs001934.v1.p1 [https://www.ncbi.nlm.nih.gov/projects/gap/cgi-bin/study.cgi?study_id=phs001934.v1.p1]). The source data underlying Figures 1c, 2a–c, 2e, 3a-d, 4a-e, 5a,b,d, and 6a-c and Supplementary Figures 1c, g, 2b-e, 4a, c, d, g, 5a-c, 6a-b, 7a-c, 8a-c are provided as a Source Data file.

## Code availability

The code is available at (https://github.com/UcarLab/SexDimorphismNatureCommunications). The R Shiny app is publicly shared at (https://immune-aging.jax.org/). Code for the R Shiny app is available at https://github.com/TheJacksonLaboratory/Ucar_Aging_Shiny.

